# The KN domain of KANK proteins contains separable talin-binding and intramolecular interaction modules

**DOI:** 10.64898/2026.06.09.731086

**Authors:** Rejina B. Khan, Till Kallem, Abhimanyu K. Singh, Benjamin T. Goult

## Abstract

KANK proteins link integrin adhesions to the cortical microtubule stabilising complex (CMSC) through interactions with the adhesion adaptor talin. However, how KANK proteins are regulated remains unclear. Here we show that the KN domain of KANK proteins contains separable regions that mediate talin binding and a conserved intramolecular interaction. Using fluorescence polarisation, NMR spectroscopy and structural analysis, we map an interaction between the N-terminal KN domain and the C-terminal ankyrin repeat domain and identify residues 60-68 of the KN domain as required for this intramolecular interaction. In contrast, the canonical LD motif within residues 30-60 mediates binding to talin. Deletion of residues 60-68 disrupts the intramolecular interaction while preserving talin binding, demonstrating that the KN domain contains distinct modules for talin engagement and intramolecular regulation. This regulatory architecture is conserved across the KANK family, although sequence variation modulates the strength of the intramolecular interaction. Together, these findings identify a modular organisation within the KANK KN domain that separates talin recognition from intramolecular regulation and is consistent with an autoinhibitory mechanism.

## Introduction

Integrin-mediated adhesions function as signalling platforms that coordinate interactions between the extracellular matrix and the actin and microtubule cytoskeletal systems. The adaptor protein talin plays a central role in these structures by activating integrins, linking them to the actin cytoskeleton and serving as a scaffold for the recruitment of signalling and cytoskeletal proteins [1]. Microtubules target to, and interact with, focal adhesions (FAs) via a network of scaffolding proteins that form the cortical microtubule stabilising complex (CMSC) [2–6]. The CMSC localises to FAs through a direct interaction between the KANK family proteins and the core FA component talin [2]. KANK proteins coordinate the recruitment of microtubules to adhesion sites [2], and this targeting of microtubules to the basal surface of the cell contributes to polarity and microtubule dynamics.

The KANK family of proteins consists of four members in vertebrates, KANK1-4, all of which share a similar domain organisation (Figure 1A) consisting of an N-terminal KN domain, a coiled-coil region, and a C-terminal ankyrin repeat domain [4, 5, 7]. In KANK1, the first coiled-coil region has been shown to bind to liprin-β1, while the ankyrin repeats recruit the kinesin-4 KIF21A, which inhibits polymerisation and prevents microtubule overgrowth in the vicinity of FAs (Figure 1B) [4]. The ankyrin repeats of KANK2 have also been shown to bind to KIF21A via residues which are conserved between the two isoforms [8]. The KN domain contains an invariant leucine-aspartic acid motif (LD motif) [9, 10] that binds to the R7 domain of talin [2, 11, 12]. Although the LD motif mediates binding to talin, whether other regions of the KN domain contribute to regulation of KANK function remains unclear.

**Figure 1.**
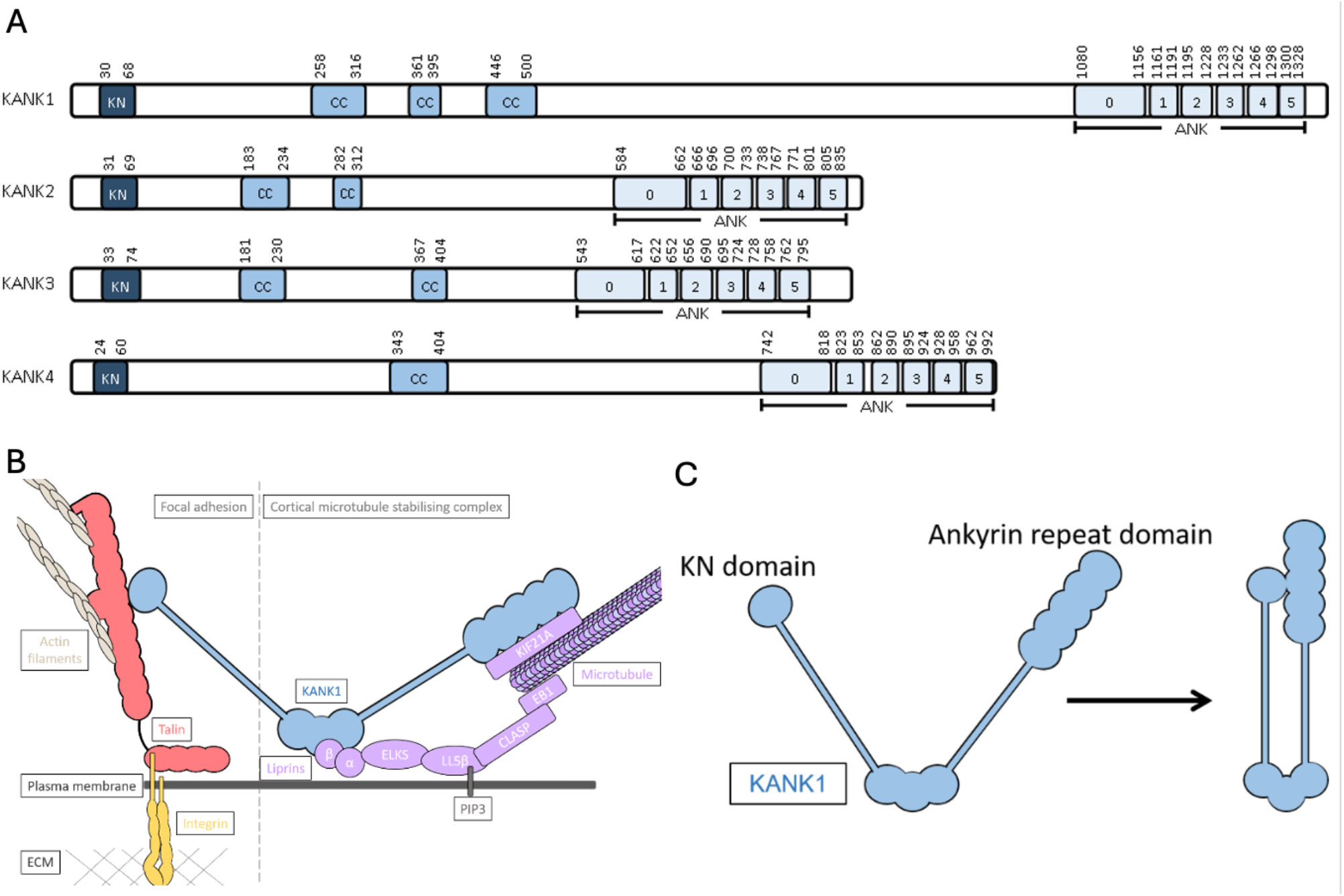
Domain organisation of KANK proteins and proposed head-tail intramolecular interaction. **(A)** Domain architecture of the four human KANK proteins. KN, KN motif; CC, coiled-coil region; ANK, ankyrin repeats. All isoforms are shown with the same horizontal scale. **(B)** KANK links integrin-mediated adhesions to the microtubule cytoskeleton. The cortical microtubule stabilising complex (CMSC, purple) assembles around the periphery of integrin adhesions through a direct interaction between the KANK KN domain and the talin rod domain R7. The kinesin KIF21A binds to the ankyrin repeats of KANK and stabilises microtubules near adhesion sites. **(C)** Schematic illustrating the proposed intramolecular interaction between the N-terminal KN domain and the C-terminal ankyrin repeat domain of KANK.

## Autoinhibition as a regulatory mechanism in adhesion proteins

Many adhesion proteins are regulated through intramolecular autoinhibition, enabling signalling complexes to assemble only when appropriate molecular cues are present [13, 14]. In these autoinhibited states, proteins adopt conformations that occlude key binding sites. For example, talin resides in the cytosol in an autoinhibited conformation in which the integrin-binding site in F3 is rendered cryptic through an interaction with the talin rod domain R9 [15, 16]. Similarly, the talin interactors DLC1 (deleted in liver cancer 1) [17] and RIAM (Rap1-interacting adapter molecule) [18] are also regulated by intramolecular head-tail interactions that partially occlude their talin-binding sites (TBS). It is likely that this mode of regulation extends to many other talin ligands.

However, despite the central role of KANK proteins in linking adhesions to the microtubule cytoskeleton, the molecular mechanisms that regulate KANK function remain poorly understood. As the talin-KANK interaction is important for microtubule targeting to adhesion structures, we hypothesised that KANK proteins might also be regulated through an intramolecular interaction between the KN domain and the ankyrin repeat domain. In this study, we identify separable regions within the KANK KN domain that mediate talin binding and intramolecular interaction, supporting a model in which KANK proteins adopt a head-tail intramolecular architecture (Figure 1C).

## Results

### The KN domain contains distinct talin-binding and ankyrin repeat-binding regions

The KN domains of all four KANK proteins have been shown to bind to the R7 domain of talin [2], whereas the C-terminal ankyrin repeats of KANK1 and KANK2 have been shown to engage the kinesin KIF21A [4]. As these interactions are important for the localisation and function of KANK proteins at adhesions, we hypothesised that KANK proteins may be regulated through an intramolecular interaction between the KN domain and the ankyrin repeats. To test this, we used an *in vitro* fluorescence polarisation (FP) assay and a series of KANK1 KN peptides (Figure 2A). A BODIPY-labelled KANK1 KN(30-68) peptide was titrated with increasing concentrations of the KANK1 ankyrin repeats. The KN peptide bound directly to the ankyrin repeats with a K_d_ of 12.9 ± 6.5 μM (Figure 2B), demonstrating an interaction between the N- and C-terminal regions of KANK1 that could support an intramolecular head-tail interaction

### Talin-1 R7R8 and KANK1 ankyrin repeats recognise distinct regions of the KANK1 KN domain

The KN domain of KANK1 has been mapped to residues 30-68 [7]. Our previous characterisation of KN domain binding to talin [2] showed that a truncated KN peptide comprising residues 30-60 (Figure 2C, green) bound talin R7 with comparable affinity to the full KN domain peptide comprising residues 30-68 (Figure 2C, red). Furthermore, the talin-KANK interaction can be disrupted by a 4A mutation that replaces the LDLD motif (residues 41-44) with AAAA (Figure 2A,C, orange) [2].

**Figure 2.**
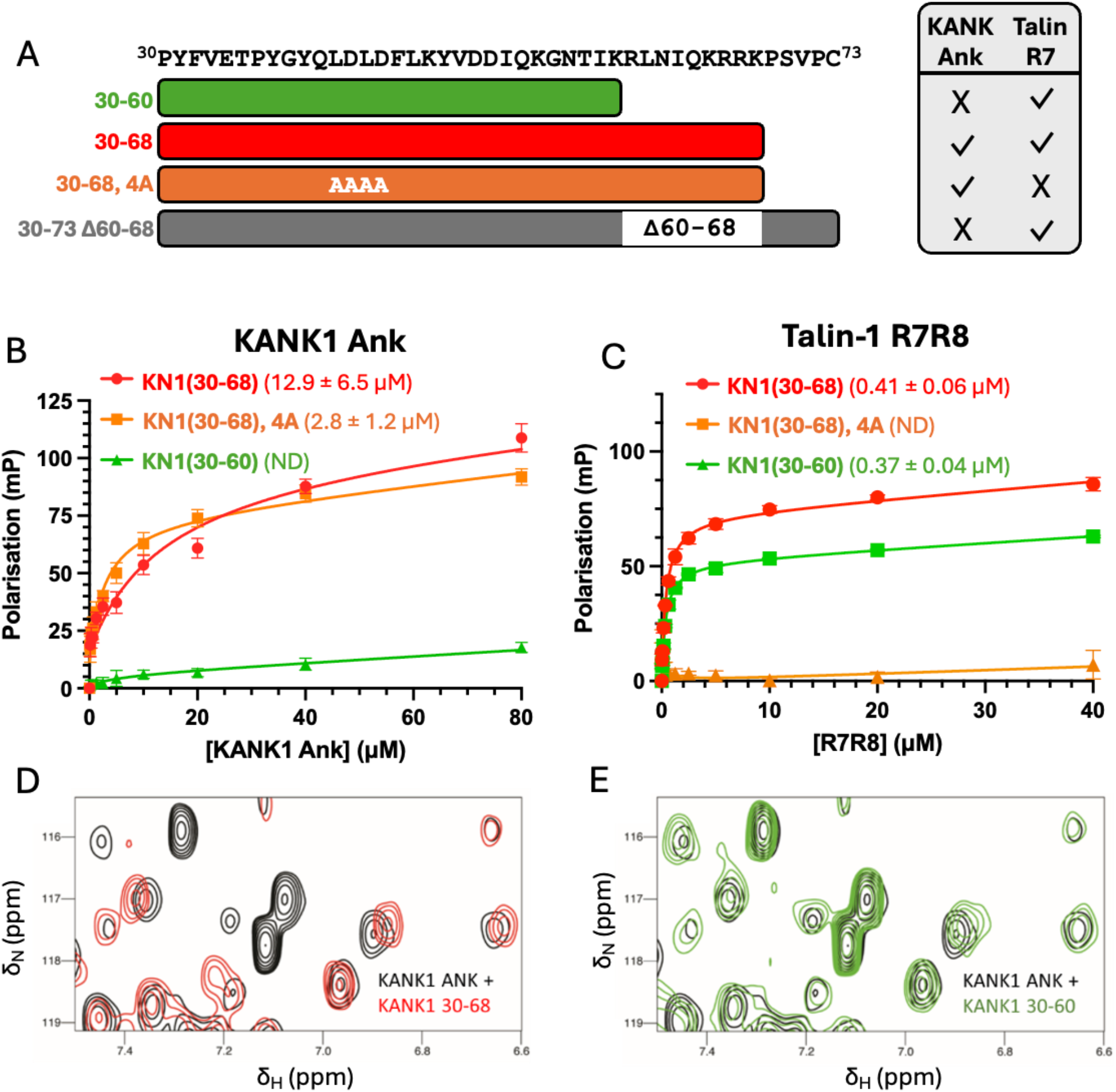
Talin-1 R7R8 and KANK1 ankyrin repeats recognise distinct regions of the KANK1 KN domain. **(A)** KANK1 KN domain peptides used in this study. **(B**,**C)** Binding of **(B)** KANK1 ankyrin repeats and **(C)** talin-1 R7R8 to BODIPY-labelled KANK1 KN peptides measured using fluorescence polarisation. Dissociation constants ± SE are indicated in the legend. All measurements were repeated in triplicate. ND, not determined. **(D**,**E)** ^1^H,^15^N TROSY NMR spectra of 100 µM KANK1 ankyrin repeats in the absence (black) or presence of 100 μM **(D)** KANK1 KN(30-68) (red) **or (E)** KANK1 KN(30-60) (green).

The KN(30-68) peptide interacted with both talin R7R8 and KANK1 ankyrin repeats, but with two striking differences in their binding requirements. First, whereas the 4A mutation abolished talin binding, it did not disrupt binding to the ankyrin repeats, with a modest increase in affinity (K_d_ = 2.8 ± 1.2 µM compared with ∼10 µM for wild-type), indicating that the LDLD motif is not required for the interaction with the ankyrin repeat domain. Second, whereas both the short and long KN peptides bound talin with comparable affinity, binding of the KN(30-60) peptide to the ankyrin repeats was not detectable under these conditions.

To further characterise the interaction, we used NMR spectroscopy to probe binding of unlabelled KANK1 KN(30-68) (Figure 2D, red) or KN(30-60) (Figure 2E, green) peptides to ^15^N-labelled KANK1 ankyrin repeats at a 1:1 molar ratio. Addition of the KN(30-68) peptide resulted in large chemical shift perturbations, consistent with the interaction observed by fluorescence polarisation. In contrast, addition of the KN(30-60) peptide produced only subtle spectral changes, further supporting a requirement for residues 60-68 in the interaction.

### Residues 60-68 in the KANK1 KN domain are required for binding to the ankyrin repeat domain

A KANK1 KN domain peptide spanning residues 30-73 but lacking residues 60-68 was synthesised. This KN(30-73Δ60-68) peptide was used in FP assays against both talin-1 R7R8 and KANK1 ankyrin repeats (Figure 3D). The deletion peptide retained high-affinity binding to talin-1 R7R8, consistent with residues 30-60 being sufficient for talin binding. In contrast, deletion of residues 60-68 abolished detectable binding to the KANK1 ankyrin repeats.

**Figure 3.**
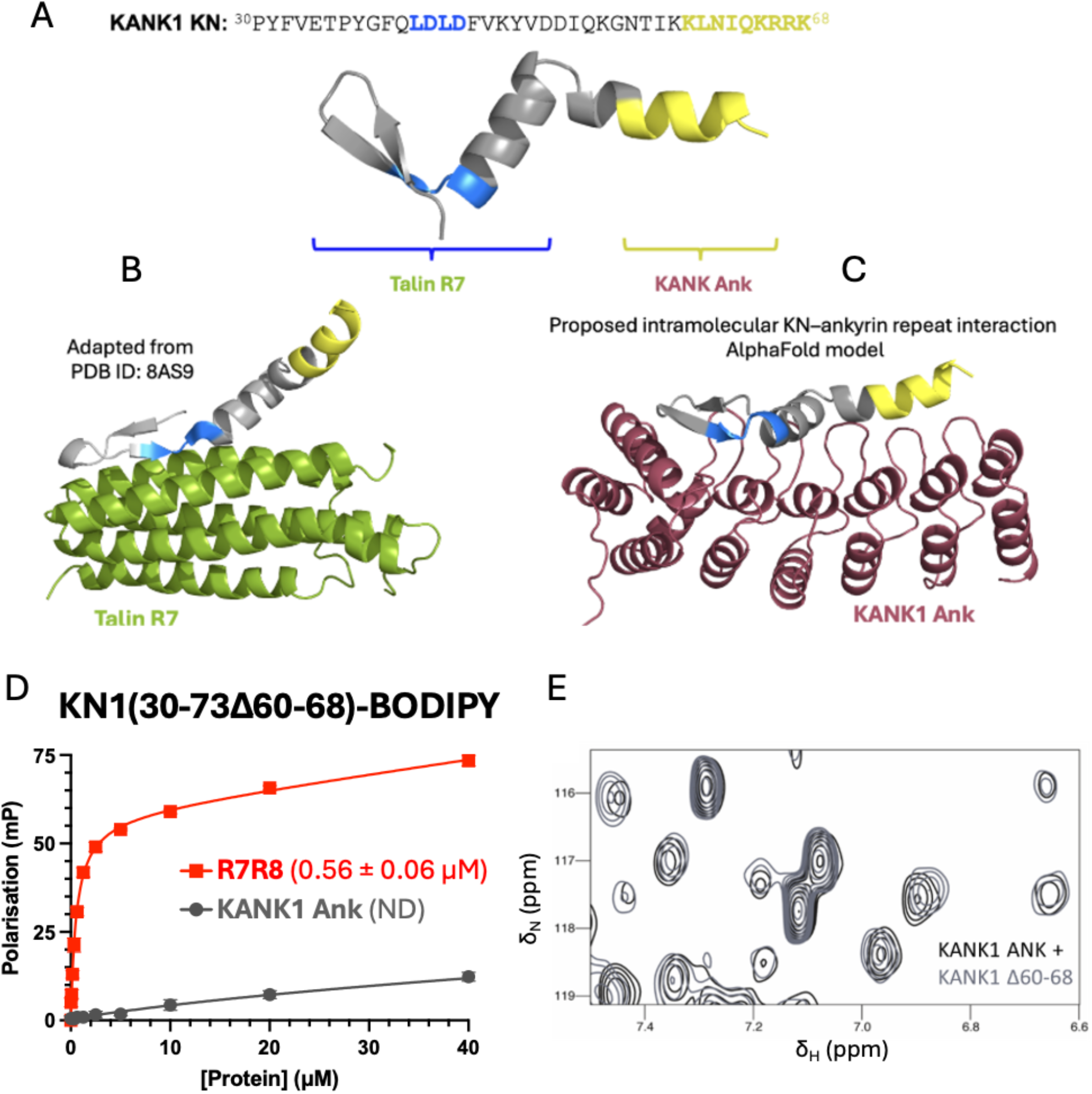
The KN domain of KANK proteins contains separable talin-binding and intramolecular interaction modules. **(A)** AlphaFold model of the KANK1 KN domain with talin-binding LDLD motif in blue and the region required for ankyrin repeat binding in yellow. **(B)** Structure of KANK1 KN domain bound to talin R7, adapted from PDB ID: 8AS9. **(C)** AlphaFold model of KANK1 KN domain bound to the KANK1 ankyrin repeats. **(D)** Binding of talin-1 R7R8 and KANK1 ankyrin repeats to BODIPY-labelled KANK1 KN (30-73Δ60-68) was measured using fluorescence polarisation. Dissociation constants ± SE are indicated in the legend. All measurements were repeated in triplicate. ND, not determined. **(E)** ^1^H,^15^N TROSY NMR spectra of 100 μM KANK1 ankyrin repeats in the absence (black) or presence (grey) of 100 µM KANK1 KN(30-73Δ60-68).

NMR spectra likewise showed very few peak shifts when the KN(30-73Δ60-68) peptide was added to ^15^N-labelled KANK1 ankyrin repeats (Figure 3E), consistent with the loss of detectable binding observed by fluorescence polarisation.

To visualise the organisation of these two binding regions, we examined an AlphaFold model of the KANK1 KN domain (Figure 3A). The talin-binding LDLD motif and the residues required for ankyrin repeat binding occupy distinct regions of the KN sequence. The experimentally determined structure of the KN domain bound to talin R7 (PDB ID: 8AS9) illustrates the interaction mediated by the LDLD motif (Figure 3B), whereas an AlphaFold model of the KN-ankyrin repeat complex provides a possible structural arrangement for the interaction identified here (Figure 3C). Together, these models illustrate how the KN domain could support two distinct interactions through separable binding regions.

### Intramolecular KN-ankyrin repeat interactions are conserved across the KANK family

We next sought to test whether KN-ankyrin repeat interactions were conserved across the KANK family. We generated peptides corresponding to the KN domains of all four KANK proteins (Figure 4A). Sequence alignment revealed that basic residues within the region corresponding to KANK1 residues 60-68 are broadly conserved, although KANK3 contains a three-residue insertion (Gly-Pro-Pro) within this region (Figure 4A). Using fluorescence polarisation assays, we found that the ankyrin repeat domains of all four KANK proteins bound their respective KN domain peptides (Figure 4B). KANK1, KANK2 and KANK3 exhibited broadly comparable affinities (K_d_ ∼12-25 µM), whereas KANK4 showed weaker binding for which a reliable K_d_ could not be determined under these conditions.

**Figure 4.**
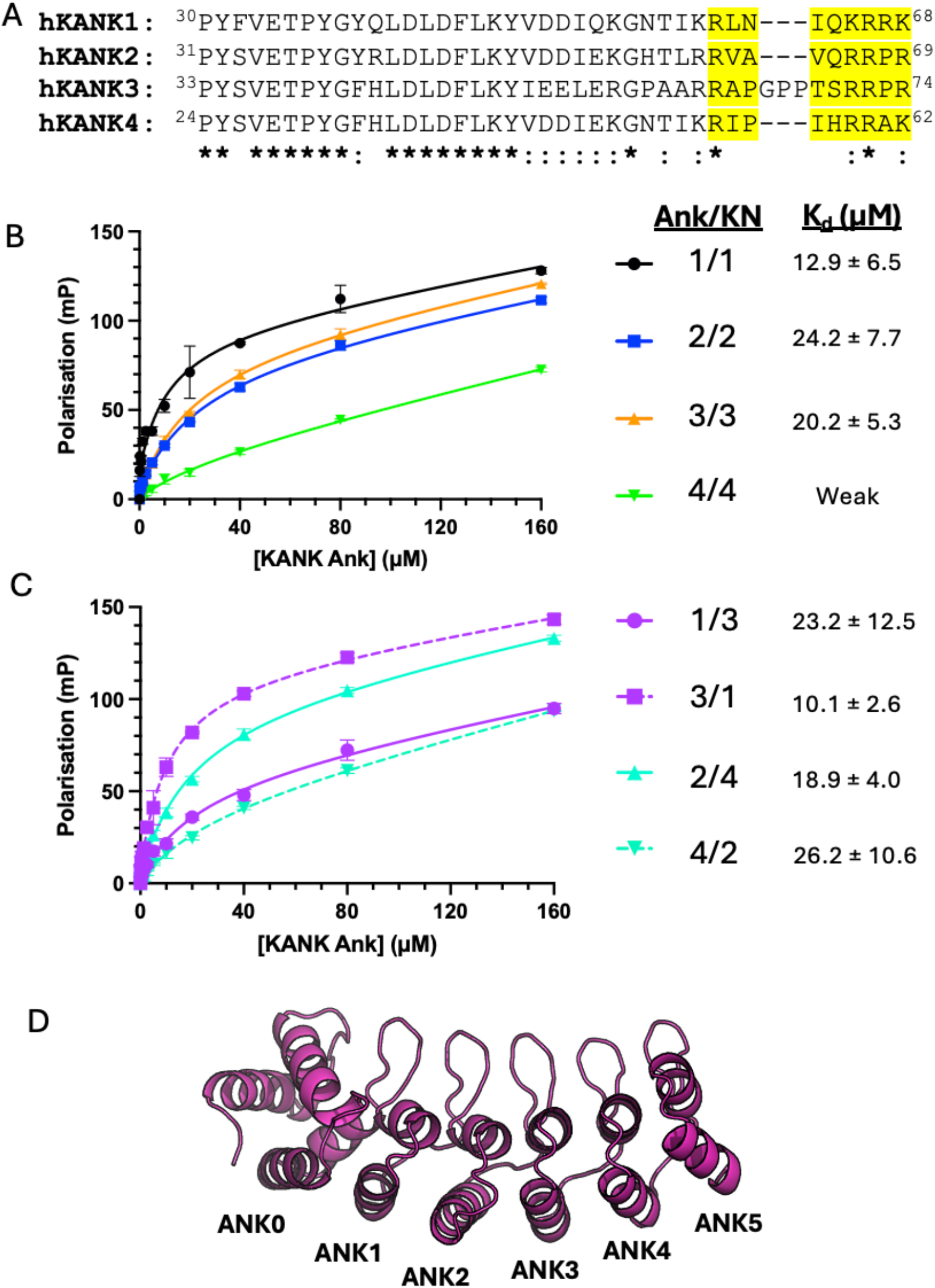
Conservation of KN-ankyrin repeat interactions across the KANK family. **(A)** Sequence alignment of the KN domains of human KANK1-4. The region corresponding to KANK1 residues 60-68 is highlighted in yellow. KANK3 contains an additional three-residue GPP insertion within this region. **(B)** Binding of the ankyrin repeat domains of KANK1-4 to their respective KN peptides measured using fluorescence polarisation. **(C)** Cross-isoform binding experiments between the KN and ankyrin repeat domains of KANK1 and KANK3, and KANK2 and KANK4, measured using fluorescence polarisation.

Dissociation constants ± SE are indicated. All measurements were performed in triplicate. ND, not determined. **(D)** Crystal structure of the KANK3 ankyrin repeat domain (PDB ID: 6TLH).

To investigate the conservation of this interaction further, we exchanged KN and ankyrin repeat domains between KANK1 and KANK3, and between KANK2 and KANK4 (Figure 4C). These experiments demonstrated that KN and ankyrin repeat domains from different KANK family members could interact, supporting conservation of the underlying recognition mechanism. The domain exchanges also suggested that differences in binding affinity may be influenced by the ankyrin repeat domain, although the effects were modest.

### The structure of the KANK3 ankyrin repeats resembles

KANK1-2 ankyrin repeats To investigate the structural conservation of the ankyrin repeat domains across the KANK family, we solved the crystal structure of the KANK3 ankyrin repeats (Figure 4D; PDB ID: 6TLH). The structure revealed high structural similarity to the ankyrin repeat domains of KANK1 and KANK2 (PDB IDs: 5YBJ [19] and 4HBD, respectively), consistent with the conserved KN-ankyrin repeat interactions observed in our binding assays. The structure was solved by molecular replacement using the KANK2 ankyrin repeat structure as a search model. Data collection and refinement statistics are summarised in Table 1

### A disease-causing mutation in KANK2 ankyrin repeats has only minor effects on the intramolecular KN-ankyrin repeat interaction

The KANK2 A670V mutation has been identified as a cause of palmoplantar keratoderma and woolly hair [20]. As this mutation is located within the ankyrin repeat domain, we investigated whether it might disrupt the KN-ankyrin repeat interaction and thereby contribute to the disease phenotype. We generated the A670V mutant by site-directed mutagenesis and used fluorescence polarisation to compare binding of wild-type and mutant KANK2 ankyrin repeats to the KANK2 KN domain (Figure 5A). The mutation had only a negligible effect on KN-ankyrin repeat binding.

**Table 1.** Crystallographic data collection and refinement statistics.

| Data collection | KANK3 ankyrin repeats | KANK2 ankyrin repeats<br>(A670V mutant) |
| --- | --- | --- |
| Synchrotron and Beamline | Diamond I04-1 | Diamond I04-1 |
| Space group | C2 | C2 |
| Molecule / a.s.u. | 2 | 1 |
| Cell dimensions |  |  |
| *a*, *b*, *c* (Å) | 149.43, 45.77, 69.59 | 95.77, 45.65, 51.07 |
| $\alpha$, $\beta$, $\gamma$ (°) | 90, 95.06, 90 | 90, 100.38, 90 |
| Resolution (Å) | 69.32 – 1.80<br>(1.84 – 1.80)* | 47.10 – 1.50<br>(1.52 – 1.50)* |
| *R*<sub>merge</sub> | 0.062 (0.776) | 0.056 (2.239) |
| *I* / σ*I* | 10.5 (1.5) | 10.3 (0.5) |
| CC(1/2) | 0.994 (0.811) | 0.999 (0.329) |
| Completeness (%) | 97.4 (96.7) | 99.2 (99.9) |
| Redundancy | 3.5 (3.7) | 3.4 (3.2) |
| Refinement |  |  |
| Resolution (Å) | 1.80 | 1.50 |
| No. reflections | 40455 (2956) | 32838 (2070) |
| *R*<sub>work</sub> / *R*<sub>free</sub> | 0.19/0.23 | 0.17/0.22 |
| No. atoms |  |  |
| Protein | 3547 | 1884 |
| Water | 370 | 186 |
| *B*-factors (Å<sup>2</sup>) |  |  |
| Protein | 33.53 | 28.21 |
| Water | 38.36 | 41.55 |
| R.m.s. deviations |  |  |
| Bond lengths (Å) | 0.012 | 0.009 |
| Bond angles (°) | 1.529 | 1.565 |
| Ramachandran plot |  |  |
| Favoured/outlier (%) | 98.74/0 | 98.33/0 |
| Rotamer |  |  |
| Favoured/poor (%) | 93.53/2.43 | 95.71/0.95 |
| MolProbity scores |  |  |
| Protein geometry | 1.36 (97<sup>th</sup>)<sup>^</sup> | 1.47 (87<sup>th</sup>)<sup>^</sup> |
| Clash score all atoms | 2.81 (99<sup>th</sup>)<sup>^</sup> | 8.8 (68<sup>th</sup>)<sup>^</sup> |
| PDB accession no. | 6TLH | 6TMD |

Data collected from a single crystal.
\*Values in parentheses are for highest-resolution shell.
<sup>^</sup>Values in parentheses indicate percentile scores as determined by MolProbity.

**Figure 5.**
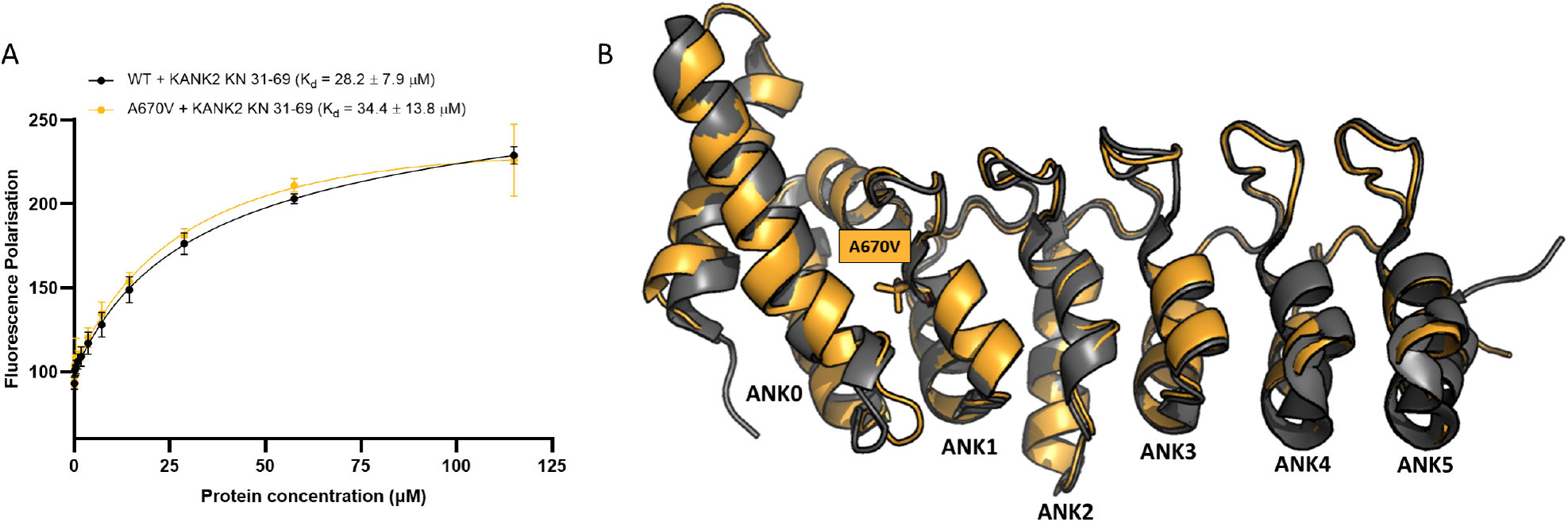
Comparison of wild-type and A670V mutant KANK2 ankyrin repeats. **(A)** Binding of wild-type and A670V KANK2 ankyrin repeats to the KANK2 KN peptide measured using fluorescence polarisation. Dissociation constants ± SE are indicated in the legend. All measurements were performed in triplicate. **(B)** Crystal structure of KANK2 A670V ankyrin repeats (yellow, PDB ID: 6TMD) superimposed on the wild-type KANK2 ankyrin repeat structure (grey, PDB ID: 4HBD), revealing high structural similarity (RMSD = 0.601 Å over 1509 atoms).

To assess the structural consequences of the mutation, we solved the crystal structure of the KANK2 A670V ankyrin repeat domain (PDB ID: 6TMD). Crystallisation trials were performed both in the presence and absence of KN peptides; however, the resulting crystals contained only the ankyrin repeat domains. Comparison with the wild-type structure (PDB ID: 4HBD) revealed only subtle structural differences (Figure 5B), consistent with the minimal effect of the mutation on KN binding.

## Discussion

Here we identify a conserved interaction between the N-terminal KN domain and C-terminal ankyrin repeat domain of KANK proteins, consistent with a potential intramolecular autoinhibitory mechanism. Mapping of this interaction identifies a short region within the KN domain (residues 60-68 in KANK1) that is required for binding to the ankyrin repeats, distinct from the canonical talin-binding LD motif. This organisation suggests a mechanism by which KANK proteins could regulate their recruitment to adhesion complexes through intramolecular interactions that influence accessibility of the talin-binding region.

### The KN domain comprises two functionally distinct regions

The KN domain was originally identified as a unifying feature of the KANK family of proteins [7], comprising around 39 residues (residues 30-68 of KANK1) at the N-terminus. In our previous study, we demonstrated that the KN domain is a talin-binding site, with residues 30-60 being sufficient for maximal talin binding [2], whereas residues 61-68 play a negligible role in direct talin binding. Here, we show that residues 60-68 are required for binding to the C-terminal ankyrin repeat domain. The interaction is conserved across all four KANK family members, with KANK1, KANK2 and KANK3 displaying comparable affinities, while KANK4 exhibits weaker binding.

Furthermore, the ability of KN and ankyrin repeat domains from different KANK proteins to interact suggests that the underlying recognition mechanism is conserved despite sequence variation between family members.

### Functional implications of the KANK intramolecular interaction

A KN peptide spanning residues 30-68 binds both talin R7 and the KANK ankyrin repeat domain. Deletion of residues 60-68 (30-73Δ60-68 in KANK1) preserves talin binding but abolishes detectable interaction with the ankyrin repeats, whereas mutation of the LDLD motif (residues 41-44) disrupts talin binding while preserving the KN-ankyrin repeat interaction. Thus, the talin-binding and ankyrin repeat-binding regions within the KN domain can be perturbed independently. These observations support a model in which KANK proteins adopt a head-tail intramolecular interaction that may regulate accessibility of the talin-binding region.

In cells, a KANK1 4A mutant exhibits a strong phenotype, with loss of localisation of KANK and the CMSC to adhesions, and a resulting loss of microtubule targeting to adhesions [2]. In contrast, despite considerable efforts to identify a comparable cellular phenotype associated with disruption of the KN-ankyrin repeat interaction, its functional consequences remain unclear. KANK proteins have been implicated in a range of cellular processes beyond adhesion organisation, including the regulation of microtubule dynamics, cell migration and disease-associated phenotypes, suggesting that mechanisms controlling KANK activity may have broader biological consequences [21]. It is possible that additional cellular or mechanical cues regulate formation of the intramolecular interaction, or that KANK proteins exist predominantly in a more open conformation under standard cell culture conditions. It remains possible that disruption of the intramolecular interaction may have more pronounced effects in alternative contexts such as 3D culture or during development.

Intramolecular head-tail interactions are a recurring feature of adhesion proteins and their associated signalling complexes. Talin itself is maintained in an autoinhibited conformation through interactions between the FERM domain and rod domains [15, 16], while several talin-binding partners, including RIAM [18], DLC1 [22] and vinculin [23], also use intramolecular interactions to regulate accessibility of talin-binding motifs. Our findings suggest that KANK proteins adopt a similar modular architecture, in which talin recognition and KN-ankyrin repeat interactions are mediated by separable regions within the KN domain. More broadly, the regulated exposure of short linear interaction motifs may represent a common organisational principle within adhesion signalling networks that coordinate actin and microtubule cytoskeletal systems.

## Materials and Methods

### Expression and purification of recombinant polypeptides

Human KANK2 ankyrin repeats (residues 583-832) and murine KANK3 ankyrin repeats (residues 524-773) were obtained as synthetic genes (GeneArt). Human KANK1 ankyrin repeats (residues 1078-1328) were amplified by PCR from human KANK1 cDNA. Murine KANK4 ankyrin repeats (residues 755-1002) were amplified by PCR from murine KANK4 cDNA, a generous gift from Reinhard Fässler. All constructs were cloned into the pET151-TOPO expression vector. Murine talin-1 (TLN1) R7R8 (residues 1357-1653) was expressed and purified as described previously [24, 25].

All proteins were expressed in *E. coli* BL21(DE3) in LB media for unlabelled protein or, for ^15^N-labelled samples for NMR, in 2M9 minimal medium using ^15^NH^4^Cl as the sole nitrogen source. His-tagged proteins were purified by nickel-affinity chromatography, with subsequent cleavage of the His-tag using TEV protease. Further purification was carried out via anion-exchange chromatography. Extinction coefficients at 280 nm (calculated using ProtParam) were used to determine the concentrations of each protein.

### Peptides

KANK KN domain peptides were synthesised by GLBiochem (China). Peptides contained a non-native terminal cysteine to allow fluorophore coupling; for most peptides the cysteine was located at the C-terminus, whereas the KANK2 peptide contained an N-terminal cysteine.

mKANK1 KN(30-60) PYFVETPYGFQLDLDFVKYVDDIQKGNTIKK-C

mKANK1 KN(30-68) PYFVETPYGFQLDLDFVKYVDDIQKGNTIKKLNIQKRRK-C

mKANK1 KN(30-68, 4A) PYFVETPYGFQAAAAFVKYVDDIQKGNTIKKLNIQKRRK-C hKANK1 KN(30-73Δ60-68) PYFVETPYGYQLDLDFLKYVDDIQKGNTIKPSVP-C

mKANK2 KN(31-69) C-PYSVETPYGYRLDLDFLKYVDDIEKGHTLRRVAVQRRPR

mKANK3 KN(32-73) PYSVETPYGFHLDLDFLKYVEEIERGPASRRTPGPPHARRPR-C

mKANK4 KN(24-65) PYSVETPYGFHLDLDFLKYVDDIEKGHTIKRIPIHRRAKQAK-C

### Fluorescence polarisation assays

Peptides were coupled to BODIPY TMR dye (Invitrogen) via the terminal cysteine using previously described methods [2]. Briefly, 1-5 mM peptide stock solutions were prepared for each peptide in PBS (137 mM NaCl, 2.7 mM KCl, 10.0 mM Na^2^HPO^4^, 1.8 mM KH^2^PO^4^), 100 µg ml−^1^ TCEP and 0.05% Triton X-100. 100 µM peptide in PBS pH 7.4 supplemented with 0.05% v/v Triton X-100 and 100 µM TCEP was reacted with a 5-fold molar excess of dye for 2 h at room temperature. The reaction was quenched by addition of 5 mM TCEP. Uncoupled dye was removed using a PD-10 column (GE Healthcare).

A fixed concentration of 0.5 μM labelled peptide was titrated with increasing concentrations of protein to a final volume of 100 µl using PBS. Titrations were performed in triplicate. Fluorescence polarisation was measured at 25°C using excitation at 544 nm and emission at 590 nm on a Spark Multimode plate reader (Tecan). Data were analysed by non-linear curve fitting using a one-site plus non-specific binding model in GraphPad Prism.

### NMR spectroscopy

Proteins for NMR titrations were prepared in 20 mM phosphate buffer (pH 6.5), 50 mM NaCl, 2 mM DTT and 5% v/v D^2^O. ^1^H,^15^N-TROSY experiments were carried out at 298 K using a Bruker AVANCE III 600 MHz spectrometer equipped with a CryoProbe. Data were processed using TopSpin and analysed using CcpNmr Analysis [26].

### X-ray crystallography

Crystallisation trials for KANK2 ankyrin repeats A670V mutant and KANK3 ankyrin repeats were carried out at 21°C by hanging drop vapour diffusion with a 1:1 protein-to-reservoir solution ratio. Crystals of the KANK2 ankyrin repeats A670V mutant were obtained in 0.2 M ammonium acetate, 0.1 M Bis-Tris (pH 5.7) and 31% (w/v) PEG 3350. Crystals of KANK3 ankyrin repeats were obtained in 0.2 M ammonium formate and 20% (w/v) PEG 3350. Crystals were cryoprotected in the same solution supplemented with 20% v/v glycerol prior to vitrification in liquid nitrogen. Diffraction datasets were collected at 100 K on beamline I04-1 at Diamond Light Source (Didcot, UK) using a Pilatus3 6M-F detector (Dectris, Baden, Switzerland). All crystallographic data was processed using autoPROC [27], which incorporates XDS [28], AIMLESS [29] and TRUNCATE [30] for data integration, scaling and merging. The structures were determined using molecular replacement carried out with PHASER [31] employing the published KANK2 ankyrin repeats structure (PDB ID: 4HBD) as search template. Iterative rounds of real and reciprocal space refinement were performed with COOT [32] and REFMAC [33] respectively. Figure preparation was carried out with PyMOL (Schrödinger LLC, Cambridge MA, USA). Data collection and refinement statistics are summarised in Table 1. Coordinates and structure factors for the KANK2 ankyrin repeats A670V mutant and KANK3 ankyrin repeats have been deposited in the Protein Data Bank under accession codes 6TMD and 6TLH.

## Acknowledgements

We thank David Critchley and Neil Ball for critical reading of the manuscript and Gary Thompson for assistance with the NMR experiments. We acknowledge Diamond Light Source for access to beamline I04-1. B.T.G. was supported by BBSRC grants (BB/N007336/1 and BB/S007245/1), a HFSP grant (RGP00001/2016) and a British Heart Foundation grant (SP/F/23/150045). R.B.K. was supported by a University of Kent studentship.

## Data Availability

Atomic coordinates and structure factors have been deposited in the Protein Data Bank under accession codes 6TLH and 6TMD. Plasmids generated in this study have been deposited in Addgene and are available to the research community (https://www.addgene.org/Ben_Goult/).

